# N-Glycosylation is a Potent Regulator of Prion Protein Neurotoxicity

**DOI:** 10.1101/2023.02.25.530047

**Authors:** Kevin M. Schilling, Pooja Jorwal, Natalia C. Ubilla-Rodriguez, Tufa E. Assafa, Jean R. P. Gatdula, David A. Harris, Glenn L. Millhauser

## Abstract

The C-terminal domain of cellular prion protein (PrP^C^) contains two N-linked glycosylation sites, the occupancy of which impacts disease pathology. In this study, we demonstrate that glycans at these sites are required to maintain an intramolecular interaction with the N-terminal domain, mediated through a previously identified copper-histidine tether, which suppresses the neurotoxic activity of PrP^C^. NMR and EPR spectroscopy demonstrate that the glycans refine the structure of the protein’s interdomain interaction. Using whole-cell patch-clamp electrophysiology, we further show that cultured cells expressing PrP molecules with mutated glycosylation sites display large, spontaneous inward currents, a correlate of PrP-induced neurotoxicity. Our findings establish a structural basis for the role of N-linked glycans in maintaining a non-toxic, physiological fold of PrP^C^.

## Introduction

Prion diseases, or Transmissible Spongiform Encephalopathies (TSEs), are a class of fatal, infectious neurodegenerative illnesses caused by the refolding of the endogenous cellular prion protein (PrP^C^) into a toxic isoform (PrP^Sc^) (1, 2). These diseases, which are closely related to other protein aggregation disorders, including Alzheimer’s and Parkinson’s diseases, are a threat to cattle, cervids and humans. Biosynthetically mature murine PrP^C^ is composed of 208 amino acids (residues 23-230), and is post-translationally modified with a C-terminal glycophosphatidylinositol (GPI) anchor and two Asn-linked glycans at residues 180 and 196. The N-terminal segment (residues 23-125, after removal of the signal peptide) exhibits a high degree of flexibility (3). Within this segment is an octapeptide repeat (OR) domain, residues 59-90, composed of the sequence (PHGGXWGQ)_4_ in mouse PrP (where X is Gly in repeats one and four, Ser in repeats two and three), which binds the divalent ions Cu^2+^ and Zn^2+^ (4-7). The C-terminal domain (residues 126-208), which contains the glycan attachment sites, folds to a characteristic structure composed of three α-helices, numbered one through three, and two anti-parallel β-strands flanking helix 1 (8, 9).

The N-terminal domain of PrP can interact with the lipid bilayer or with other membrane proteins to cause several toxic activities, including membrane leakage, spontaneous ionic currents, and neuronal death (10-14). It has been hypothesized that these activites may be responsible for the neurotoxic effects of prions during the disease process. The activity of the N-terminal, “toxic effector domain” is normally suppressed by an intramolecular interaction with the globular C-terminal domain (11, 15). Our research has demonstrated that copper binding to the OR segment promotes a stable contact between the two PrP domains that is essential for this regulatory interaction (11, 15, 16).

Extensive mutagenesis, combined with biophysical and electrophysiological experiments provide a detailed structural picture of how the two domains of PrP interact (15). Specifically, copper drives the *cis* interdomain interaction through joint histidine coordination, in which His residues are contributed by each of the two domains (15). The C-terminal histidines (H139 and H176) are presented on a surface largely composed of α-helices two and three (Figure 1). The N-terminal histidines are primarily those in the ORs. Conserved, C-terminal acidic residues (D177, E195, E199, E210), which lie on the same protein surface as the coordinating His residues, further facilitate this interaction. Familial mutations that reduce the negative charge of this patch have been linked to inherited prion diseases (17). These include the highly penetrant E200K mutation (E199K murine sequence), which causes Creutzfeldt-Jakob disease (CJD), and D178N (D177N murine), which causes Fatal Familial Insomnia (FFI) or CJD, depending on polymorphic residue 129 (M or V, respectively).

**Figure 1.**
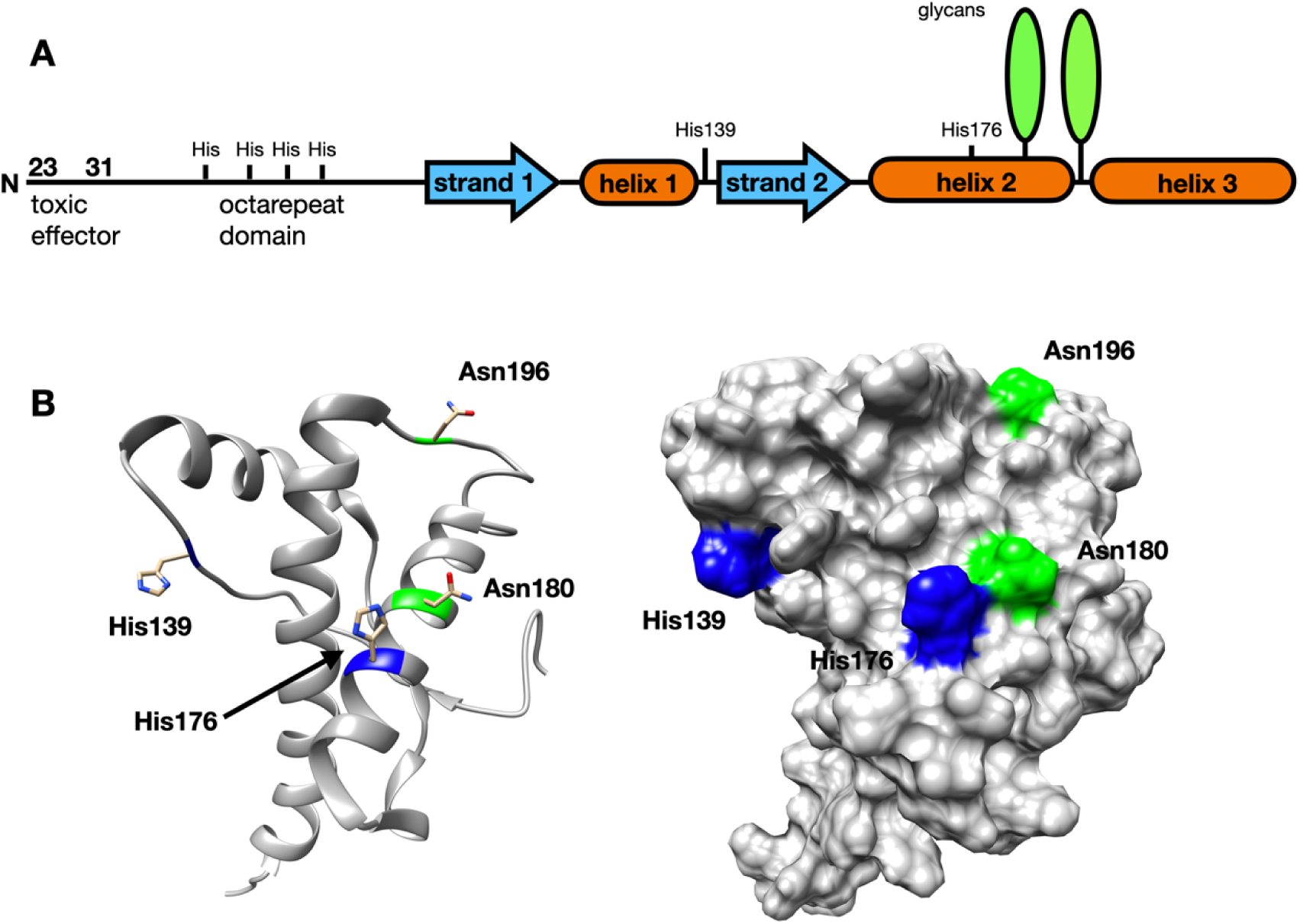
**A)** Linear diagram of the full-length PrP^C^. **B)** Ribbon and surface diagrams of the C-terminal domain of the prion protein. Note that the glycans are on the same C-terminal surface as the two histidines implicated copper coordination. Graphics use PDB: 1XYX.

PrP^C^’s N-linked glycosylation sites are at asparagines 180 and 196, proximal to the location of many of its C-terminal pathological mutations (18). The protein exists *in vivo* as a mixture of unglycosylated, monoglycosylated, and diglycosylated forms. The glycans vary in their molecular composition and size. Glycosylation patterns vary between prion strains, which are characterized by differences in incubation time, neuropathology, and clinical symptoms (19-21). For example, the PrP^Sc^ found in two human prion diseases, familial CJD caused by V180I, and variably protease-sensitive prionopathy, is devoid of glycosylation at N181(22).

Glycosylation also affects other properties of PrP^C^ and PrP^Sc^. Glycosylation partially inhibits the aggregation of PrP^C^ into infectious PrP^Sc^, decreasing its toxicity (23, 24). Glycosylation status also modulates cross species transmission – the presence of a glycan at N196 prevents transmission of sporadic human CJD to mice (23). The presence of glycans also modulates the interaction between PrP^C^ and cofactors such as glycosaminoglycans (GAGs). GAGs accelerate prion disease by enhancing the conversion of PrP^C^ into PrP^Sc^ and promoting incorporation of PrP^Sc^ into plaques; glycans reduce PrP binding to GAGs (23, 25). The terminal saccharide in the prion protein’s glycans is often sialic acid, which carries a negative charge at physiological pH. This may lead to electrostatic repulsion between copies of PrP, which could explain why sialylation slows the rate of PrP^Sc^ amplification (26-29). Prion strains show selective, strain-specific recruitment of PrP^C^ sialoglycoforms (30, 31). Lastly, the ratios of PrP^C^ glycoforms differ across brain regions, which may partially explain why prion strains have preferences for the regions that they infect (17, 32-34). In summary, the glycosylation status of the prion protein is intricately linked to many aspects of disease status, but the relationship is complicated and poorly understood.

The glycan linked to Asn180 is at a surface site near the middle of helix two, while the glycan at Asn196 extends from a loop connecting helices two and three (Figure 1). These sites are proximal to the C-terminal PrP^C^ surface containing the His residues that coordinate copper and are key to the *cis* regulatory interaction. Consequently, we deemed it essential to test whether glycosylation modulates C-terminal regulation of the N-terminal toxic effector domain. This is furter motivated by the recognition that certain prion diseases produce PrP^Sc^ with no glycosylation at Asn180, a glycosylation site that is spatially adjacent by one helix turn from His176 (Figure 1).

In this paper, we apply both Electron Paramagnetic Resonance (EPR) and Nuclear Magnetic Resonance (NMR) to artificially glycosylated, ^15^N labeled, PrP^C^ to investigate the effects of the glycans on neuroprotective self-regulatory, *cis* interaction. Using EPR, we show that glycans on PrP^C^ do not significantly inhibit or alter its copper coordination environment. NMR experiments demonstrate that the glycans refine the *cis* interaction, localizing the copper interaction surface to a more focused area on the C-terminus of the protein. Cell trafficking assays show that unglycosylated PrP^C^ is exported quantitatively to the cell surface. We showed previously that mutation of the C-terminal domain His residues impairs the observed *cis* interaction in unglycosylated PrP^C^; new data presented here shows that this effect is reversed by glycosylation. Lastly, we find that unglycosylated PrP^C^ causes cultured cells to produce spontaneous inward cationic currents, a proxy for prion protein induced toxicity. Surprisingly, the magnitude of these currents is similar to that observed for the highly toxic deletion mutant, ΔCR-PrP^C^. Taken together, these results suggest that the glycans on the surface of the C-terminal domain of the prion protein participate in maintaining a neuroprotective inter-domain interaction, working in tandem with a previously discovered copper-histidine tether.

## Results

### Glycans on the C-terminus of PrP^C^ do not significantly inhibit its copper binding

Bacterially expressed proteins lack glycans, while PrP from mammalian sources is heterogeneously glycosylated and cannot be easily isotopically labeled for structural studies. In order to create ^15^N labeled, homogenously glycosylated PrP in bacteria, we combined the genetic incorporation of the unnatural amino acid *p*-acetyl-phenylalanine with aminooxy functionalized monosaccharides, coupling them together to form an oxime linkage, as described previously (35, 36). The monosaccharides were then enzymatically extended into trisaccharides using glycosyltransferases to yield the Sial-Gal-GlcNAc trisaccharide moiety (Figure 2).

**Figure 2.**
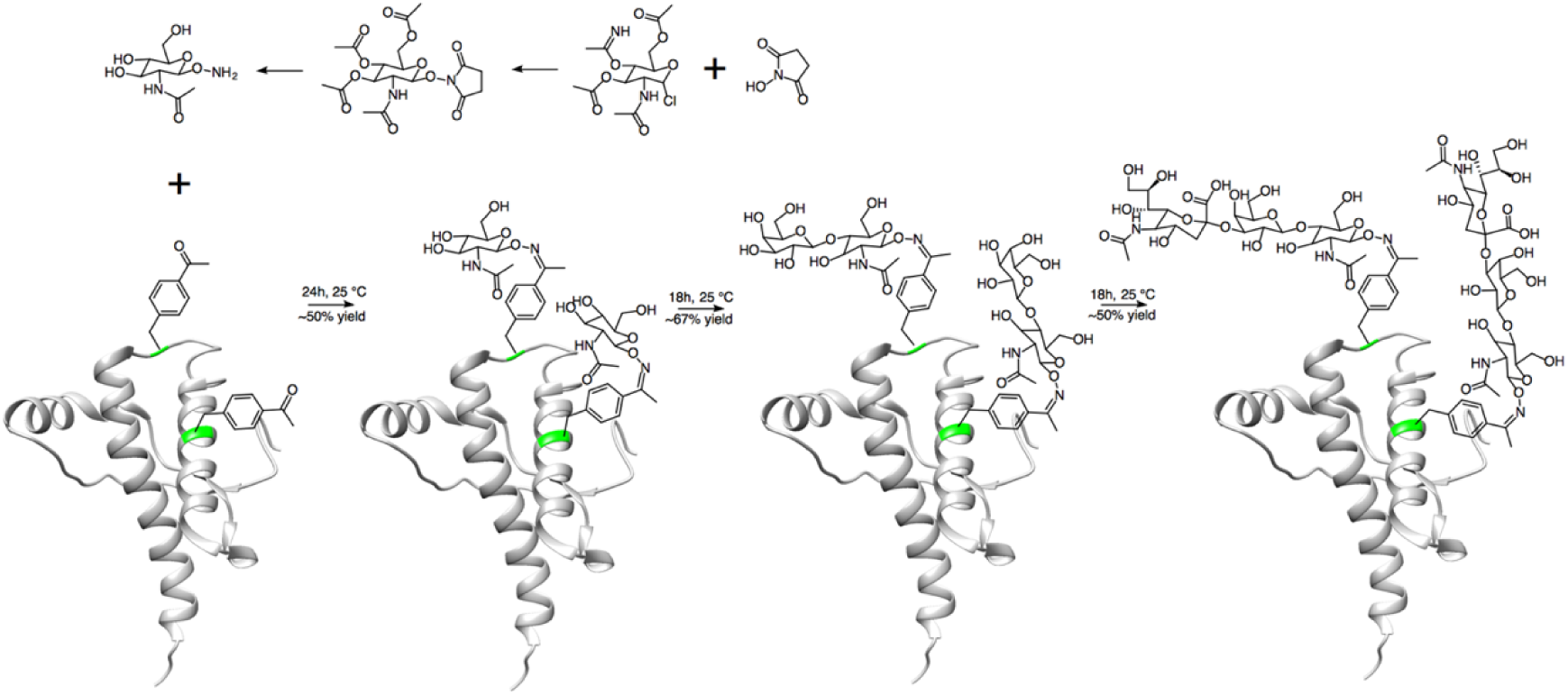
Synthesis scheme for glycosylated PrP. Plotted on PDB: 1XYX. (Adapted from Schilling et al. *J. Org. Chem*. 85:1687-90, 2020)

Continuous wave EPR was used to probe the coordination environment of the copper bound to unglycosylated and artificially glycosylated PrP^C^ at pH 6.0. Experiments were performed with uniformly ^15^N (I = ½) labeled PrP^C^, which gives better resolution of nitrogen superhyperfine coupling than protein containing the more abundant ^14^N (I = 1) isotope. Samples contained 1:1 Cu^2+^:protein, which favors so-called component 3 copper coordination, previously characterized as arising from interaction with three or four His imidazole side chains (37). Unglycosylated and glycosylated samples give nearly equivalent Cu^2+^ EPR spectra with no evidence of unbound, aquo copper (Figure 3). The parallel regions of the spectra, between 2600G and 3100G, are an exact match with g_||_ = 2.255 and A_||_ = 570 MHz, as determined from simulations (Figure 3). The values are consistent with component 3, multi-His coordination (37). The perpendicular regions match in multiplicity and field, with a slight difference in line broadening. Simulations show that these spectra are consistent with four equatorial nitrogen ligands (Figure 3). We previously demonstrated that this coordination environment arises from three OR His residues and a His on the C-terminal regulatory surface (15). Overall, these data show that glycans on the C-terminus of PrP^C^ do not inhibit or significantly alter its copper coordination modes.

**Figure 3.**
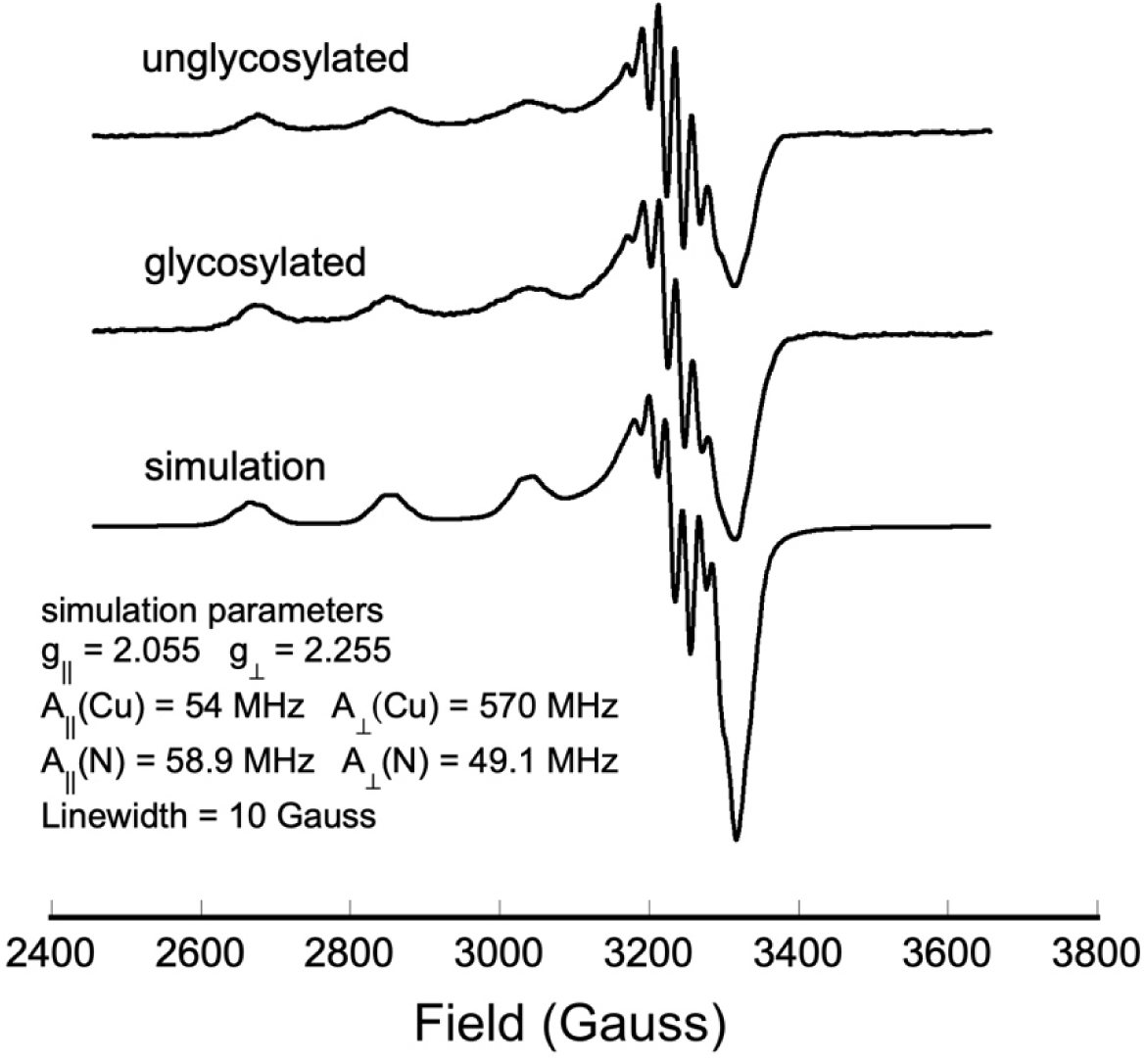
Continuous wave EPR spectra of Cu^2+^ complexed with WT unglycosylated and artificially glycosylated PrP^C^, along with a spectral simulation (parameters in the inset).

### Glycans on the C-terminus of the prion protein refine its *cis* interaction

To evaluate the respective C-terminal interaction surfaces, we employed ^1^H-^15^N HSQC NMR analysis on glycosylated and unglycosylated PrP^C^ at pH 6. Divalent copper (d^9^ electron configuration) possesses an unpaired electron that reduces the NMR signal intensity of nearby nuclei, in a distance-dependent manner, through paramagnetic relaxation enhancement (PRE). Intensities are measured from the 2D ^1^H-^15^N HSQC spectra and used to annotate residues on the PrP^C^ C-terminal domain. For clarity, only residues from 120 and beyond are displayed. Multiple representations of the data are shown in Figure 4. Figures 4A and 4B are surface representations noting glycosylation sites (green) and with intensity reduction ranked as zero (gray), weak (light blue) and strong (navy blue). Figure 4C and 4D show the same data plotted on a ribbon diagram. This representation shows clearly that all affected residues are located to helices 2 and 3, consistent with our previous findings. Figures 4E and 4F provide position-dependent intensity ratios (i/i_o_), allowing a quantitative comparison between the unglycosylated and glycosylated proteins.

**Figure 4.**
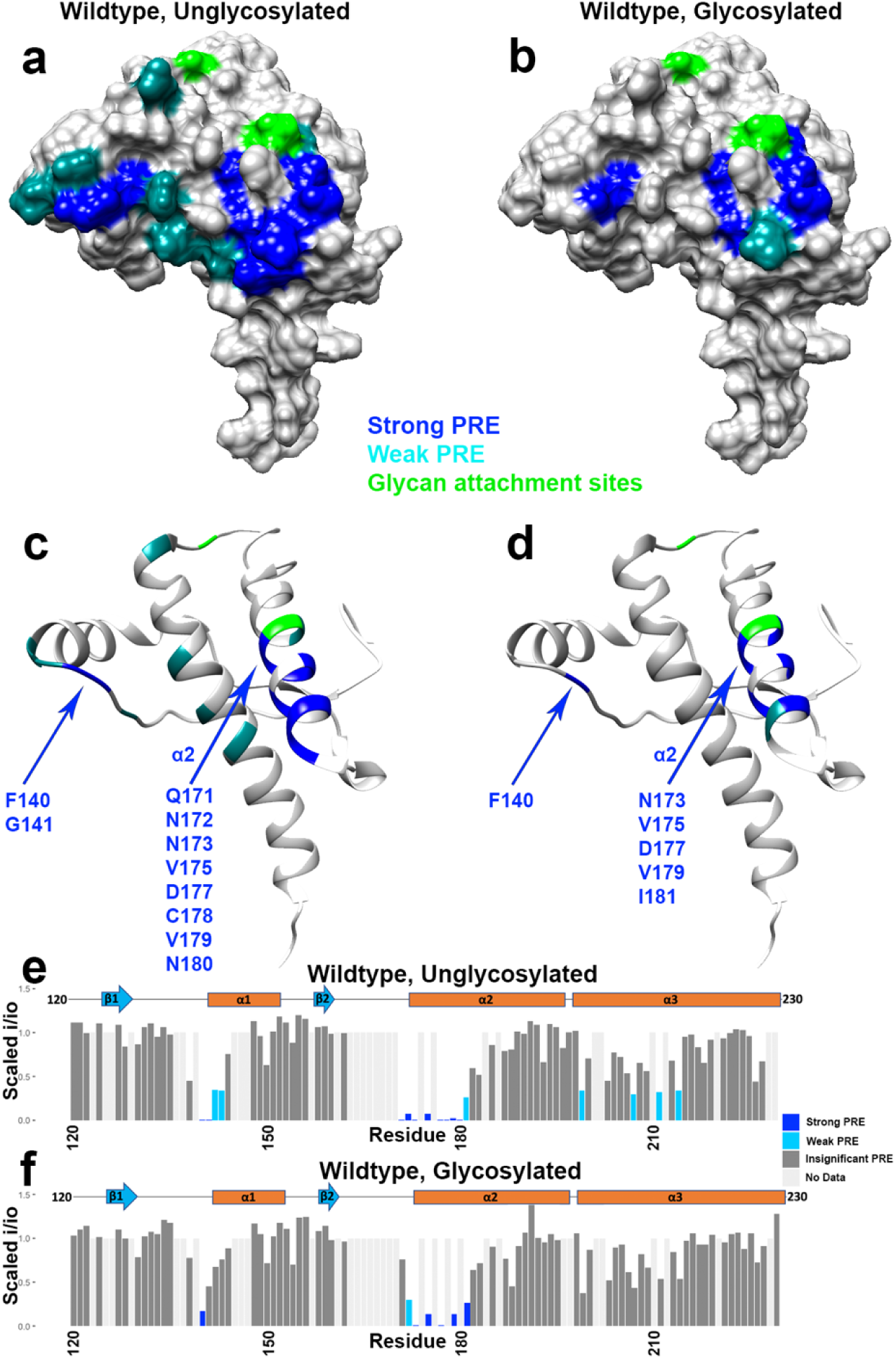
Surface representations, ribbon plots, and bar plots showing areas of wild-type PrP^C^ engaged in the *cis* interaction, with and without glycans, as measured by the intensity reduction of NMR cross peaks. Plotted on PDB: 1XYX.

We observed that the glycosylated prion protein displayed broadened signals on the C-terminal surface, like the unglycosylated protein, but that this broadening was concentrated to fewer residues and a smaller surface area (Figure 4). Specifically, while the unglycosylated protein had significant line broadening around the areas of both His139 and His176, the broadening in the glycosylated protein was focused mainly around histidine 176. This histidine is in close proximity to the glycan four residues (one helical turn) away, attached at position 180. This suggests that the glycans are not sterically hindering the *cis* interaction but, instead, are localizing the Cu^2+^-OR segment to a more well-defined surface patch, primarily on helix 2.

### Glycans partially restore *cis* interaction lost by mutation of C-terminal histidines

Histidines at positions 139 and 176 on the C-terminal domain of PrP^C^ coordinate copper together with N-terminal OR histidines, driving a neuroprotective, *cis* interaction through a metal ion molecular tether (15). We found previously that, when these C-terminal His residues are mutated to Tyr, the interaction is compromised, resulting in a much weaker interaction through a patch of negatively charged residues on the C-terminal surface (15). In addition, electrophysiological experiments performed on N2a cells expressing this mutant PrP^C^ lacking the C-terminal His residues, showed weak spontaneous transmembrane currents, consistent with increased toxicity promoted by a poorly regulated N-terminal domain. By performing ^1^H-^15^N HSQC NMR analysis on both glycosylated and unglycosylated PrP^C^(H139Y, H176Y) at pH 6, we found that the glycans largely restore the Cu^2+^-promoted line broadening consistent with the interdomain *cis* interaction that is otherwise lost by elimination of the two His residues. Specifically, whereas the unglycosylated wild-type protein has strong copper coordination around both histidines 139 and 176, the glycosylated and histidine-mutated protein has strong copper coordination around the former location of histidine 176 only (Figure 5). This suggests that the *cis* interaction in the cellular version of the protein may be exclusively between the N-terminal histidines and histidine 176, and is assisted by the nearby glycans.

**Figure 5.**
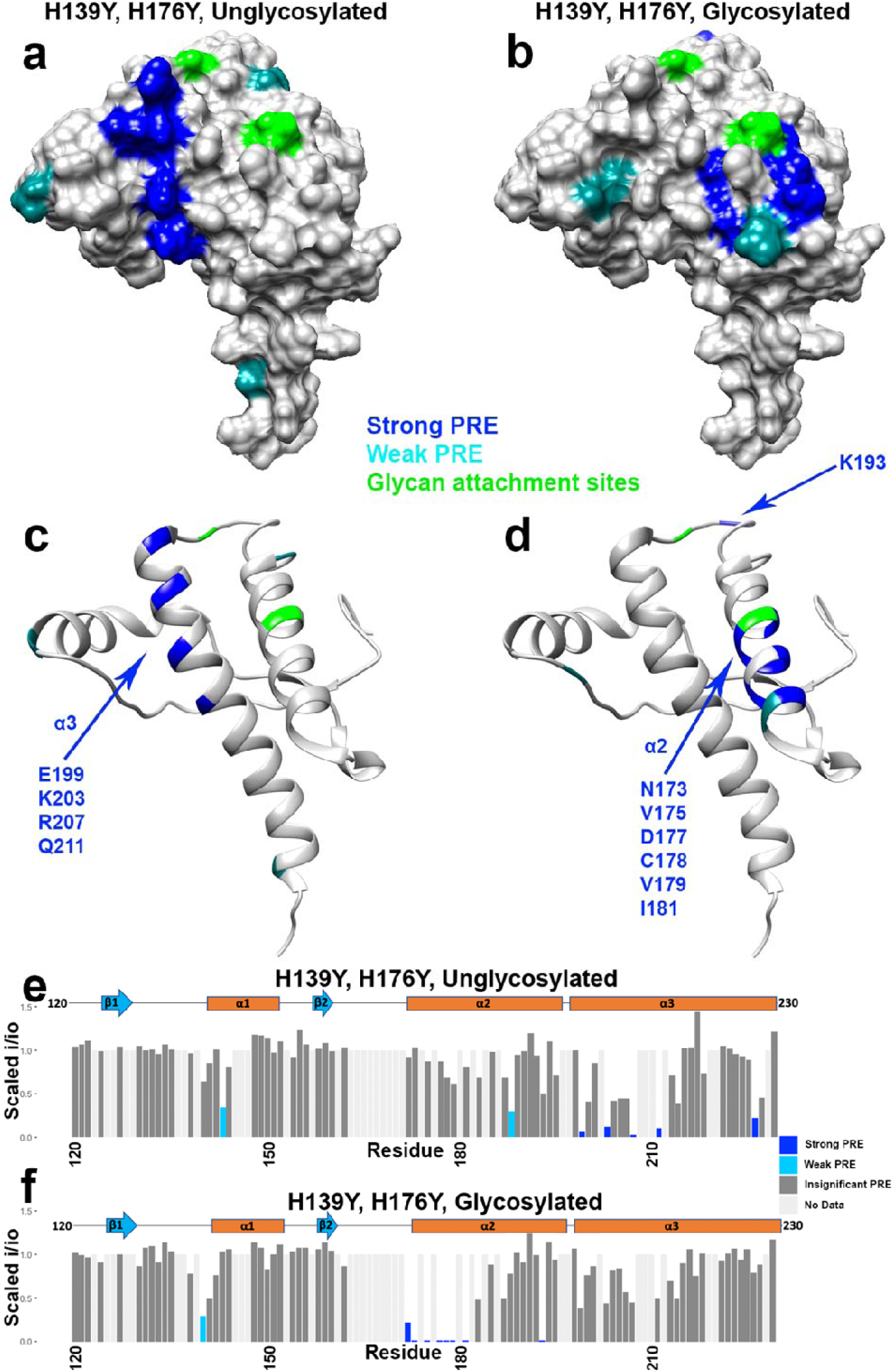
Surface representations, ribbon plots, and bar plots showing areas of H139Y, H176Y PrP^C^ engaged in the *cis* interaction, with and without glycans, as measured by the intensity reduction of NMR cross peaks. Plotted on PDB: 1XYX.

### Glycan elimination causes strong, spontaneous currents in cultured cells

Deletions in the central region of PrP, such as Δ105-125 (referred to as ΔCR), cause spontaneous, inward cationic currents in cultured cells and primary neurons (38, 39). Coexpression of wild type PrP in cultured cells suppresses these currents, mirroring the ability of coexpressed wild type PrP to suppress the neurotoxicity that accompanies these same mutations in transgenic mice (10). Antibodies that are neurotoxic when injected into the mouse brain also cause spontaneous currents in cultured cells (11). Therefore, the presence of spontaneous currents in cultured cells provides a surrogate readout of neurotoxicity caused by the prion protein. These currents are thought to be due to a weakening or abolishment of the neuroprotective *cis* interaction between the prion protein’s N and C-terminal domains, in turn allowing the polybasic N-terminal residues to form transmembrane pores (11, 40). Given our observation that glycans in the C-terminal domain enhanced the structural specificity of the *cis* interaction, we sought to test whether the absence of glycans perturbed the *cis* interaction, thereby causing spontaneous currents in cultured cells.

We created a PrP mutant, N180Q/N196Q, in which Asn residues at the two glycan attachment sites were changed to Gln residues, thereby preventing glycosylation at these sites but preserving the polar carboxamide side chain common to both Asn and Gln. In contrast to mutation of threonine residues in the N-X-T consensus sites, the N/Q mutations eliminate glycosylation but do not cause misfolding of PrP and do not alter its cellular trafficking or localization (41-44). To confirm the latter point, we expressed N180Q/N196Q PrP in N2a cells and used immunofluorescence staining to assess its cellular distribution. As shown in Figure 6, N180Q/N196Q PrP is localized to the cell surface, similar to WT PrP.

**Figure 6.**
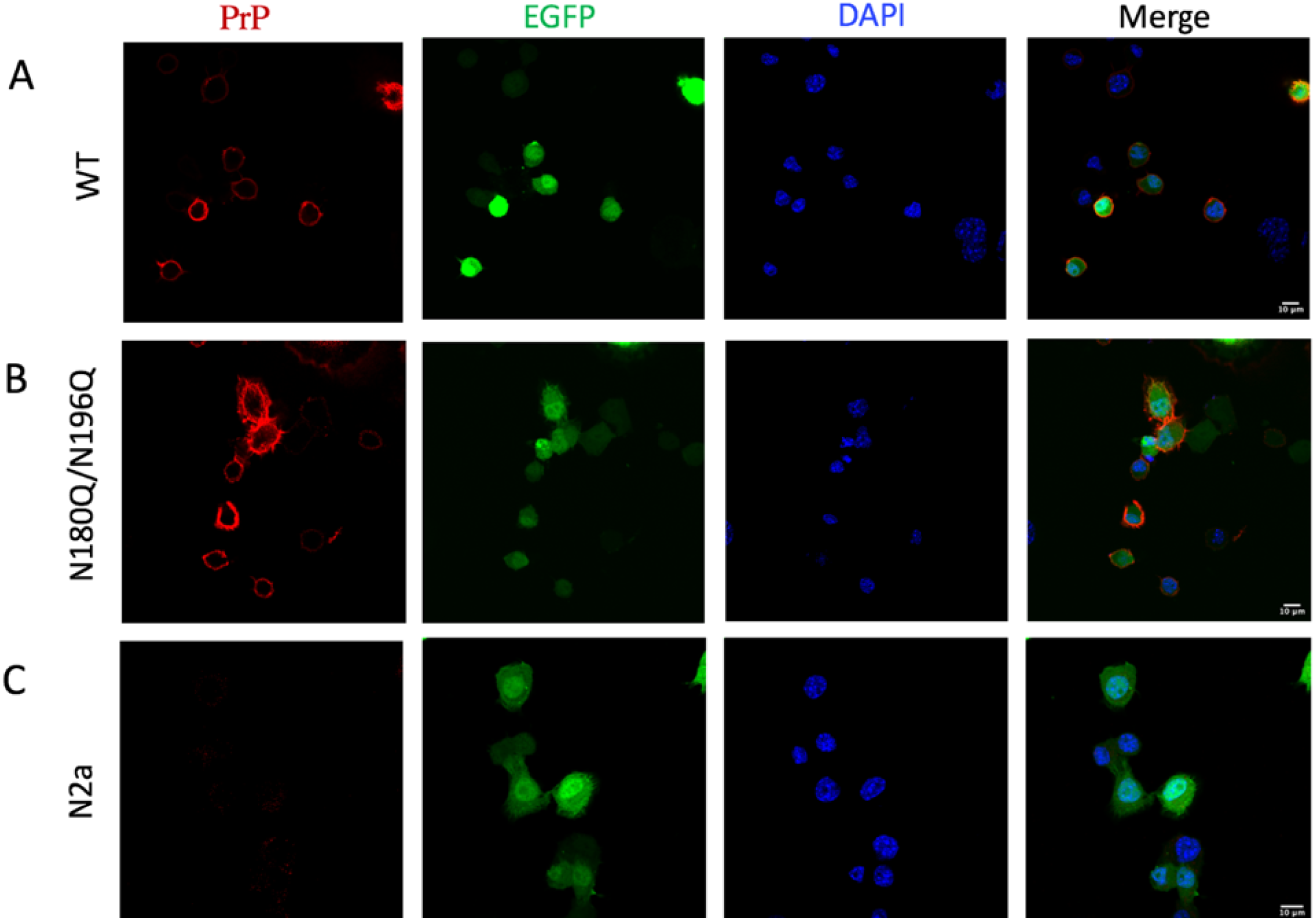
Cell surface expression of the PrP^C^ glycosylation mutant in N2a cells. N2a cells in which endogenous PrP expression was eliminated by gene editing were transfected with plasmids encoding WT PrP **(A)** or N180Q/N196Q PrP **(B)**, along with an EGFP plasmid as a transfection marker. Cells in **(C)** were transfected with the EGFP plasmid alone. Cells were then fixed without permeabilization and immunostained for PrP using D18 antibody. Images show fluorescence for PrP, EGFP, DAPI, and a merge of all three channels. The results show that N180Q/N196Q PrP is present on the cell surface, similar to WT PrP. Bar, 10 µm.

Using whole-cell patch-clamp recording with holding potentials of either −70 mV or −90 mV, we observed spontaneous inward currents for N180Q/N196Q PrP but not for the wild-type control (Figure 7; Supporting Information Figure S1). We compared these currents to ΔCR PrP as a positive control, and H139Y/H176Y PrP as a control that we previously showed produced currents at −90 mV but not at −70 mV (15). These data show that elimination of N-glycosylation causes strong spontaneous currents at −70 mV, quantitatively comparable to those observed for the highly toxic ΔCR PrP. Interestingly, we found little difference between N180Q/N196Q PrP, lacking the glycans, and the quadruple mutant N180Q/N196Q/H139Y/H176Y PrP, which lacks both glycans as well as the C-terminal His residues implicated in copper binding. Collectively, these data indicate that the C-terminal glycans play a significant role in stabilizing the interdomain *cis* interaction, thereby contributing to regulation of the toxic effector function of the N-terminal domain.

**Figure 7.**
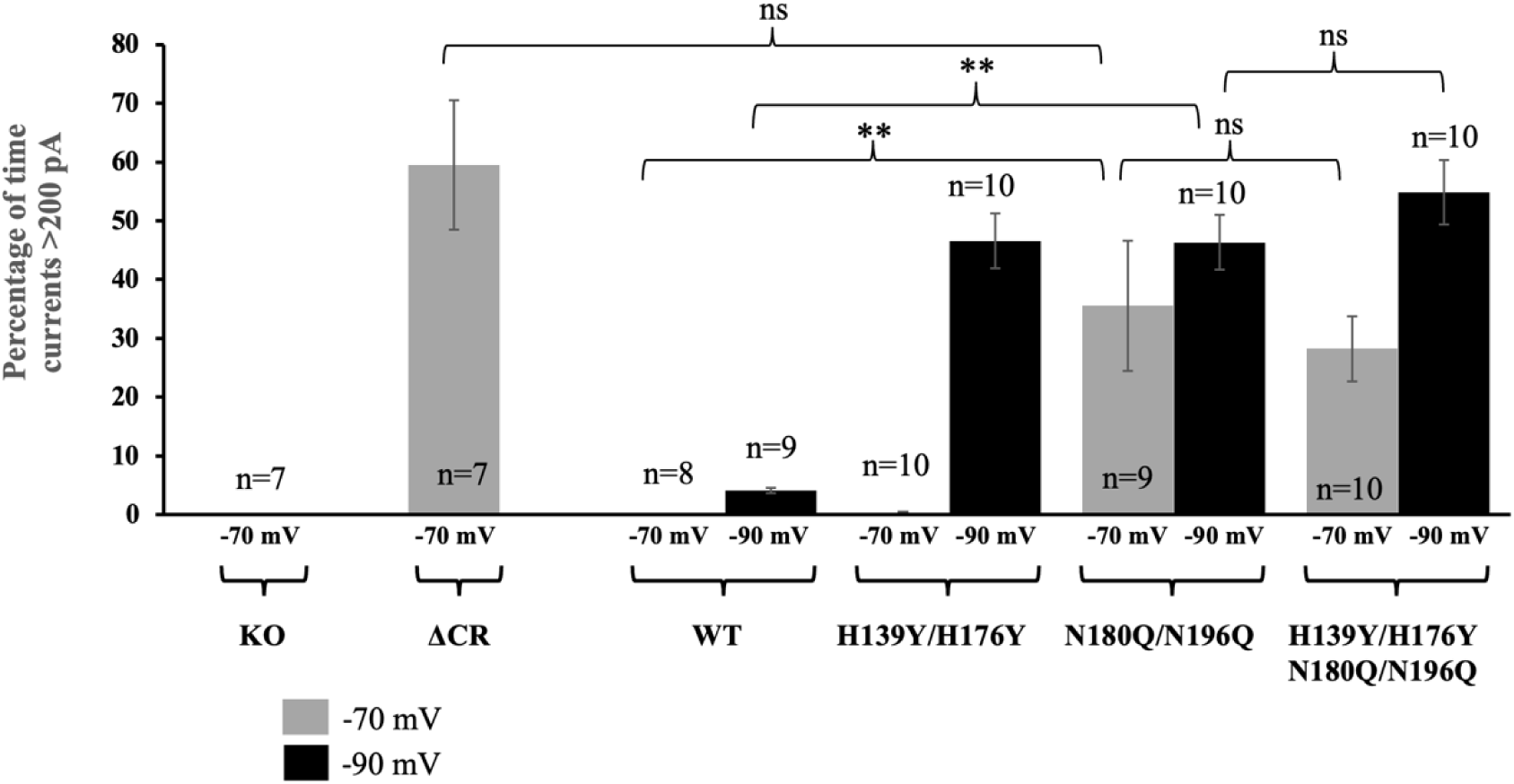
Whole-cell patch-clamp recordings from N2a cells in which endogenous PrP expression was eliminated by gene editing. Cells were untransfected (KO), or were transfected to express the following forms of PrP: Δ105-125 (ΔCR), wild type (WT), H139Y/H176Y, N180Q/N196Q and H139Y/H176Y/N180Q/N196Q. The holding potential was either −70 mV or −90 mV, as indicated. Bars show mean ± S.E.M. Differences are not significant (ns), or are significant with p<0.01 (double asterisk). Mutation of both N-linked glycosylation sites (N180Q/N196Q) significantly enhanced spontaneous inward currents at a holding poteintial of −70 mV, even in the absence of mutation of the histidine residues (H139Y/H176Y).

## Discussion

In this study, we show that glycans on the C-terminus of PrP^C^ refine the protein’s neuroprotective, copper-driven *cis* interaction, localizing domain-domain contact to a well-defined region on the C-terminal domain. We further show that the presence of the glycans partially reverses loss of this interaction caused by the mutation of C-terminal histidines to tyrosines. Lastly, we find that unglycosylated PrP^C^ causes cultured cells to produce spontaneous inward currents, a readout that serves as a measure of PrP-induced neurotoxicity. Together, these results suggest that the glycans contribute structurally to auto-regulation in the wild-type protein.

In a previous study, we found that two histidines on the C-terminus of the prion protein drive a neuroprotective *cis* interdomain interaction by tethering the C-terminus to an N-terminally bound copper ion. Here, we find that the PrP glycans also promote an N-C interaction, synergizing with the effect of His Cu coordination. A patch of negatively charged amino acids, located on the same protein surface as the histidines and glycans, is a third contributor (16, 45). More specifically, the cumulative data suggest that His176, acting through copper coordination, glycans at Asn180, and acidic residues D177, E195, E199, E210 act in concert to anchor the toxic effector N-terminal domain to its regulatory site on the C-terminal domain, as shown schematically in Figure 8.

**Figure 8.**
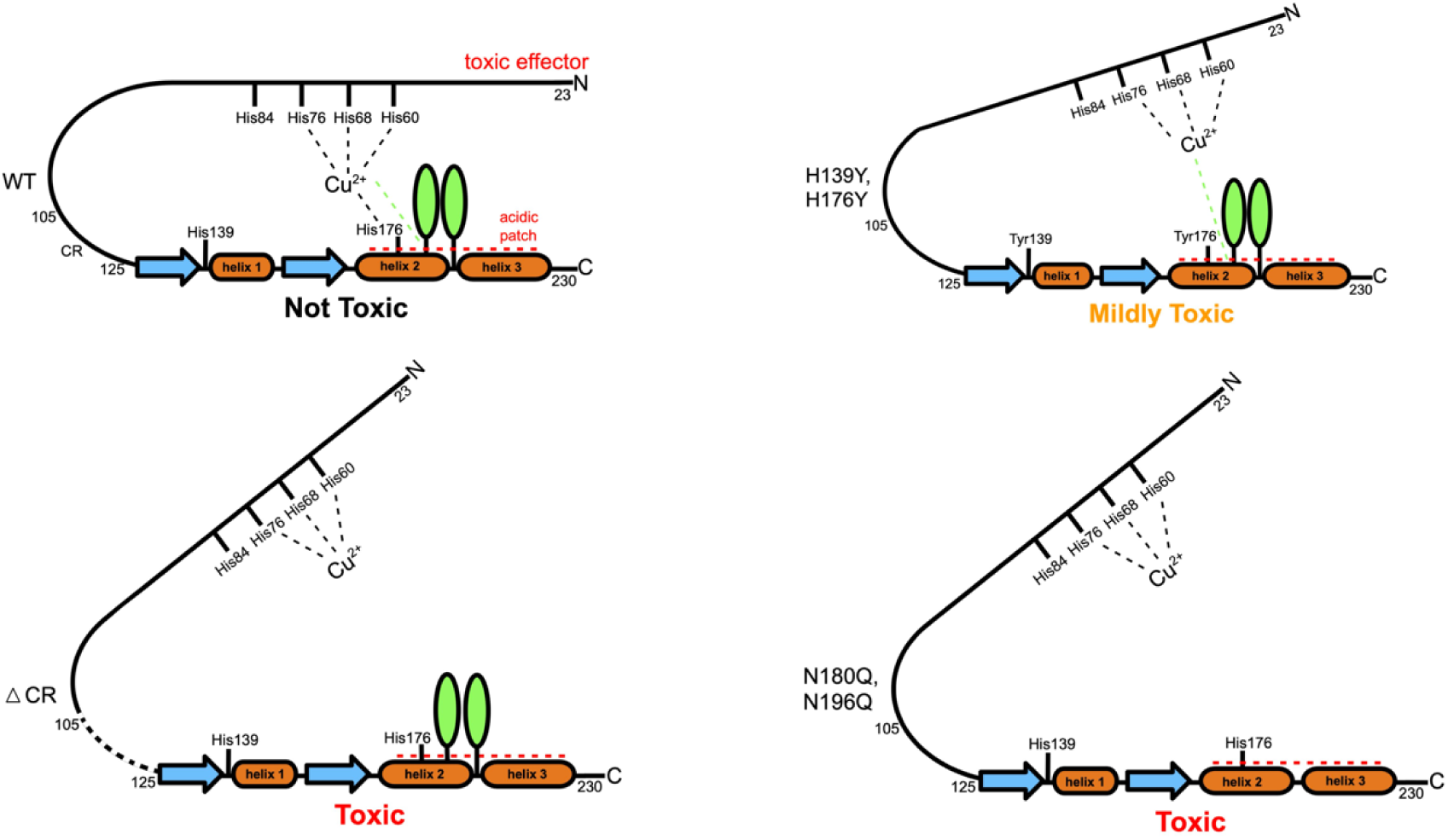
Unified model showing three contributors to the *cis* interaction in PrP^C^: the negatively charged patch produced by acidic residues in the C-terminal domain (red lines), Cu^2+^ coordinating to His residues in the OR and the regulatory C-terminal domain and glycans in the C-terminal domain (teal ovals). ΔCR PrP^C^ is the deletion mutant lacking residues 105-125.

Figure 8 summarizes a qualitative ranking among the various factors that control regulation of the N-terminal, toxic effector domain, as measured by the strength of the spontaneous currents induced by different mutants. Several familial forms of prion disease arise from mutations of acidic, C-terminal residues, including D177N and E199K (D178N and E200K in the human sequence). Previous work with the Zn^2+^-PrP^C^ complex show that these mutations weaken the metal ion-promoted cis interaction. However, these mutations do not cause spontaneous currents in cultured cells. Next, we found that elimination of the C-terminal histidines, by His→Tyr mutagenesis, results in weak spontaneous currents that are observable only at hyperpolarizing transmembrane voltages of −90 mV. This effect is categorized as being “mildly toxic” (Figure 8). Previous to this present study, extensive investigations with the deletion mutant PrP^C^(Δ105-125) (ΔCR PrP^C^), which eliminates a portion of the linker between the toxic effector domain and the regulatory domain, revealed very strong spontaneous currents (38, 39). Moreover, ΔCR PrP^C^ produces a neonatal lethal phenotype in Tg mice (10). Thus, we categorize this mutant as “toxic.” In our previous studies, no other PrP mutant produced spontaneous currents as large as those observed for ΔCR PrP^C^. Our present finding that the glycosylation mutant N180Q/N196Q produces such large, spontaneous currents is therefore unexpected, and reveals a major role of N-linked glycans in regulating the toxic activity of the N-termimnal domain.

How do the C-terminal glycans stabilize the interaction between the N- and C-terminal domains of PrP, and how does this effect synergize with that of copper coordination and binding to the acidic patch? In the absence of glycosylation, both C-terminal histidines (H139 and H176) contribute to copper coordination, as seen by enhanced PRE at the two sites. However, with glycosylation, this interaction shifts preferentially to His176, which is one helical turn away from the glycosylation site at position 180. Thus, the additional glycan moiety at position 180 may provide an enhanced surface area for interaction with the hydrophobic OR domain. Alternatively, it is well-established that protein glycosylation stabilizes protein-protein interactions through charge complementarity. For example, the tetrameric legume soybean agglutinin exhibits enhanced stability compared to other members of the same protein family by virtue of ionic contacts between its glycans and adjacent amino acid side chains (46). PrP^C^ possesses two polybasic charge clusters: one at the N-terminus (residues 23-28) and the other between the OR and globular C-terminal domains (residues 100-109). The glycans terminate with negatively charged sialic acid residues, which may interact with these positively charged clusters, thereby stabilizing the folded state. In this case, loss of the glycan chains would destabilize the folded state, freeing the N-terminal domain to assume additional conformations.

Given the strong ionic currents induced by the double glycosylation mutant (N180Q/N196Q), which are similar to those induced by the highly pathogenic ΔCR mutant (Figure 8), it might be predicted that expression of unglycosylated PrP in mice might produce a spontaneous neurodegenerative illness. To our knowledge, however, there are only two published studies featuring transgenic mice expressing PrP glycosylation mutants (47, 48), but both of these involved substitution of Ala for Thr in the N-X-T consensus site, mutations that are known to cause misfolding of PrP (41-43). These mice were analyzed for their susceptibility to prion infection, but no information was provided on the health of uninoculated mice. In light of our findings, it would be interesting to make Tg(N180Q/N196Q) mice, and determine whether they develop spontaneous neurological illness in the absence of prion infection.

## Conclusions

This work identifies a fundamentally new protective role for glycosylation of PrP^C^. Structurally, the C-terminal glycans help anchor and regulate interaction of the toxic, N-terminal effector domain with a regulatory surface on the C-terminal domain. Electrophysiological experiments demonstrated that lack of this glycan-mediated interaction leads to large transmembrane ionic currents, which could compromise cellular function. Thus, PrP^C^ glycosylation plays a crucial role in the structural and functional properties of PrP.

## Materials and Methods

### Protein Preparation

The protein used for the experiments in this study was produced using a previously published protocol (36). Briefly, plasmids containing the genes for PrP and the necessary machinery to incorporate *p*-acetyl-phenylalanine were transformed into *E*.*coli*. The bacteria was grown in minimal media with ^15^N ammonium chloride and *p*-acetyl-phenylalanine added, producing PrP with two *p*-acetyl-phenylalanine residues at the locations of the glycan attachment points. The protein was purified by nickel affinity chromatography and reverse phase high performance liquid chromatography (HPLC). The protein was allowed to react with aminooxy N-Acetylglucosamine, which attached to the two *p*-acetyl-phenylalanines through oxime linkages. Glycosyltransferases were then used to extend these sugar into trisaccharides, and the protein was once again purified by HPLC. The resulting protein was glycosylated at both residues 180 and 196 with the trisaccharide N-acetylglucosamine, galactose, sialic acid.

### Electron Paramagnetic Resonance Spectroscopy

All samples were made to pH 6.0 in 50 mM 2-(*N*-morpholino)ethanesulfonic acid (MES) buffer (Sigma), using potassium as a counterion. The protein was added to a concentration of 100 μM, and CuCl_2_ was used at 100 μM. The samples contained 30% glycerol as a cryoprotectant. X-band (9.38 GHz) continuous-wave (CW) EPR spectra were recorded on a Bruker (Billerica, MA) EleXsys E580 spectrometer equipped with a super-high Q resonator (ER4122SHQE). Cryogenic temperatures were achieved with a liquid nitrogen finger dewar and gas flow controller. The spectrometer settings were as follows: temperature = 121 K, conversion time = 41 ms, modulation amplitude = 0.5 mT, modulation frequency = 100 kHz, bridge power = 5 mW, attenuation = 23 dB.

### Nuclear Magnetic Resonance Spectroscopy

All samples were made to pH 6.0 in 10 mM 2-(*N*-morpholino)ethanesulfonic acid (MES) buffer (Sigma), using potassium as a counterion and containing 10% D_2_O. For all samples, the protein was added to a concentration of 100 μM. For samples with copper, CuCl_2_ was used at 100 μM. ^1^H-^15^N HSQC spectra were recorded at 37 ^°^C on a Bruker 800 MHz spectrometer at the UCSC NMR facility. NMR spectra were analyzed with NMR Pipe and Sparky using assignments transferred from previous experiments by visual inspection, and figures were made using Chimera, R, and python. To determine a cutoff i/i_0_ value to separate the residues involved in the *cis* interaction from the rest of the protein, we performed a kernel density estimation on the data using a Gaussian smoothing kernel. To eliminate the effects of differential unspecific peak intensity reduction across mutants, the data were scaled so that the center values of each mutant’s group of unaffected peaks were aligned. We divided the residues into three categories based on their i/i_0_ values: strongly affected (dark blue), weakly affected (light blue) and unaffected (grey). These divisions were created by using the local minimum separating the affected from unaffected residues in the wild-type protein (i/i_0_ = 0.35), and dividing the affected peaks into two groups (i/i_0_ = 0 to 0.175, and i/i_0_ = 0.175 to 0.35).

### Cell Culture

Mouse N2a cells (CCL-131, from the ATCC) were maintained in Opti-MEM Reduced Serum Medium (Thermo Fisher #31985088) supplemented with 10% (v/v) fetal bovine serum, and 1% (v/v) penicillin/streptomycin (Thermo Fisher #15140122). Cells were maintained at 37 °C in a humidified incubator containing 5% CO_2_ and were typically passaged every 4 days at a dilution of 1:5.

### Generation of PrP^-/-^ N2a Cells

Ablation of endogenous PrP expression in N2a cells was done through CRISPR-Cas9 gene editing. N2a.PrP^-/-^ clones were generated using a multi-guide sgRNA system, followed by single cell cloning by limiting dilution. Briefly, three 1.5 nmol dry sgRNAs (Synthego), UCAGUCAUCAUGGCGAACCU, GGGCCAGCAGCCAGUAGCCA, and UCAUGGCGAACCUUGGCUAC were dissolved in 15 µL of nuclease-free 1x TE buffer to make a stock solution of 100 pmol/µL of sgRNA. This was pulse vortexed for 30 seconds and incubated at room temperature for 5 minutes to fully dissolve the sgRNA. A final concentration of 30 pmol/µL of the sgRNAs and 20 pmol/µL of a Cas9 2NLS nuclease (Synthego) were introduced to 1.0 × 10^6^ cells through electroporation using Lonza’s Amaxa Cell Line Nucleofector Kit V protocol (Program T-024). Cells were plated in 6-well dishes for recovery. After 24 hrs. or until 80-90% confluence, the cells were passaged into a T-75 flask (Fisher Scientific, # FB012937) for expansion. After 3-4 days or until confluence, N2a.PrP^-/-^ cells were trypsinized, counted, and diluted to 2.0 cells/mL of medium, and 500 µL were plated in 48-well plates. The plates were monitored for 2 weeks for the presence of single-cell N2a.PrP^-/-^ clones. Selected colonies were expanded sequentially in 12-well and then 6-well plates. N2a.PrP^-/-^ clone candidates were analyzed and validated by Western blot analysis, and using Synthego’s Inference of CRISPR Edits (ICE) software. Clones that attained a knockout score of ≥ 90% in ICE and showed no trace of endogenous PrP^C^ expression on Western blots were selected and expanded further. A relatively fast-growing N2a.PrP^-/-^ clone (Clone B5) was used for all experiments.

### Whole-Cell Patch Clamp Experiments

N2a cells were maintained in DMEM supplemented with nonessential amino acids, 10% fetal bovine serum, and penicillin/streptomycin. Whole-cell patch clamp recordings were done from N2a cells 24-48 hr after transient transfection, using Lipofectamine 2000, with pEGFP-N1 (Clontech) along with pcDNA3.1 vector encoding WT or mutant PrP. The N2a cells used here had been gene-edited using CRISPR-Cas9 to disrupt the endogenous PrP gene. Recordings were done for 5 min durations using standard whole-cell patch-clamp technique (Supporting Information Figure S1). Transfected cells were recognized by green florescence. Pipettes were pulled from thick-walled borosilicate glass filament with resistance 3-5 MΩ. Experiments were conducted at room temperature with an external recording solution containing 150 mM NaCl, 4 mM KCl, 2 mM CaCl_2_, 2 mM MgCl_2_, 10 mM glucose, and 10 mM HEPES (pH 7.4 with NaOH); and an internal recording solution containing140 mM Cs-glucuronate, 5 mM CsCl, 4 mM MgATP, 1 mM Na_2_GTP, 10 mM EGTA, and 10 mM HEPES (pH 7.4 with CsOH). Current signals were acquired using a Multiclamp 700B amplifier (Molecular Devices, Sunnyvale, CA), digitized with a Digidata 1440 interface (Molecular Devices), and saved to disc for analysis with PClamp 10.7 software.

### Immunofluorescence

24-48 hrs after transfection with EGFP, WT PrP and mutant PrP plasmids, N2a cells were fixed with 4 % paraformaldehyde in PBS and treated with 0.5% BSA. PrP was detected using D18 antibody and Alexa Fluor 633-conjugated goat anti-human as secondary antibody, and nuclei were stained with DAPI. All images were acquired with a Zeiss LSM 700 confocal microscope under 63x magnification and analyzed using ImageJ.

## Acknowledgements

We gratefully acknowledge NIH grants R35 GM131781 (to G.L.M.) and R01 NS065244 (to D.A.H.) for financial support, and NIH instrumentation grants S10OD018455 and S10OD024980 for acquisition of the 800 MHz NMR spectrometer and pulsed EPR spectrometer, respectively. We also thank Drs. K. Sakamoto and S. Yokoyama at the RIKEN Yokohama branch, for providing the B-95.AAAfabR *E*.*coli* strain, and Amy Freiberg for help with the graphics.

