## Supporting Information for "N-Glycosylation is a Potent Regulator of Prion Protein Neurotoxicity"

Fig. S1

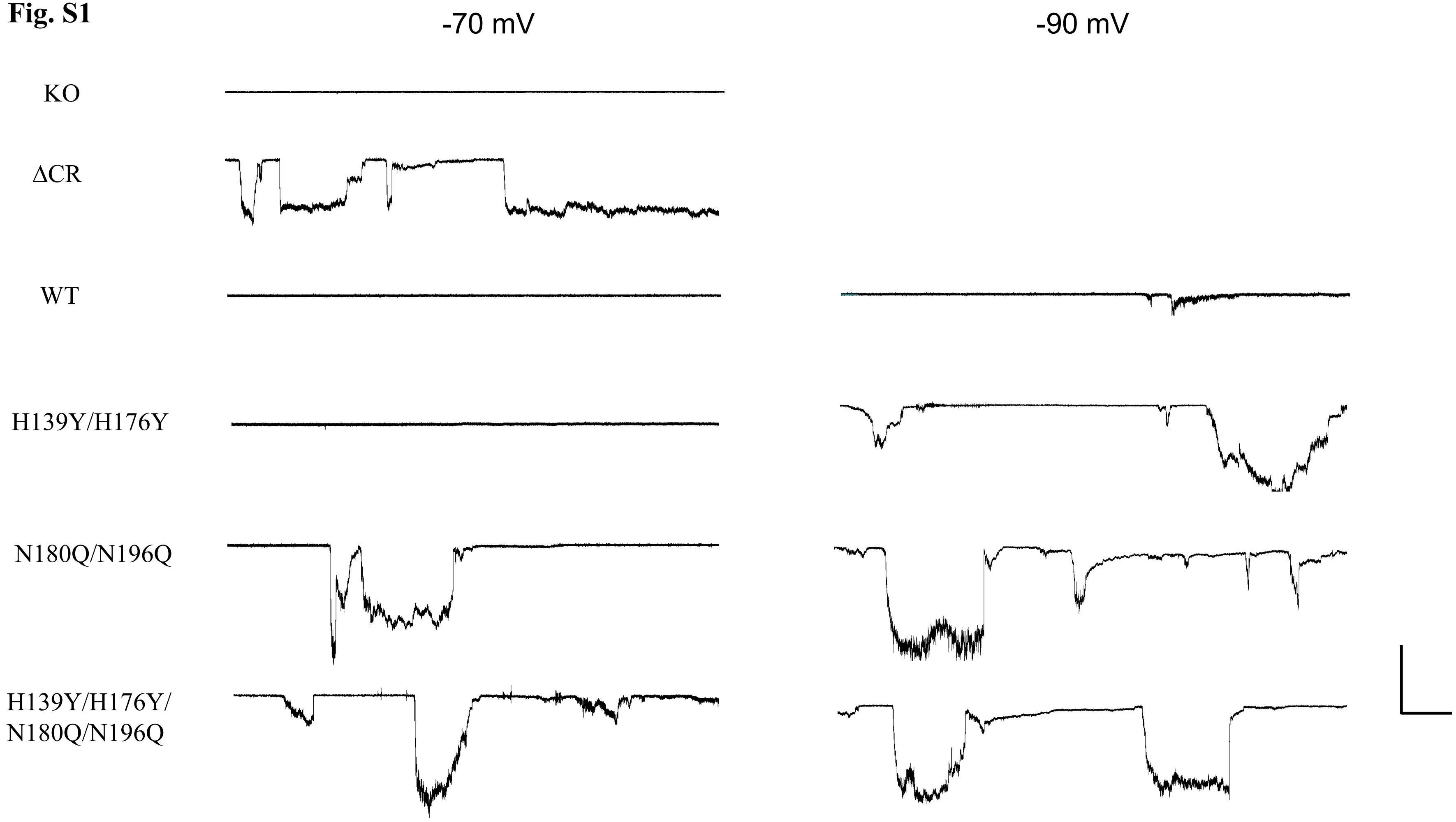


**Fig. S1** Representative traces of currents recorded from either untransfected N2a cells (KO) or N2a cells expressing following forms of PrP : ΔCR, WT, H139Y/H176Y, N180Q/N196Q and H139Y/H176Y/N180Q/N196Q. Quantification of the currents are shown in figure 7. Holding potential was either -70 mV (left) or -90 mV(right) as mentioned. Scale bar: 1 nA, 30 s.
